# Diffusion or advection? Mass transfer and diffusive boundary layer landscapes of the brown alga *Fucus vesiculosus*

**DOI:** 10.1101/094052

**Authors:** Mads Lichtenberg, Rasmus Dyrmose Nøerregaard, Michael Kühl

## Abstract

The role of hyaline hairs on the thallus of brown-algae in the genus *Fucus* is long debated and several functions have been proposed. We used a novel motorized setup for 2D-and 3D-mapping with O_2_-microsensors to investigate the spatial heterogeneity of the diffusive boundary layer (DBL) and O_2_ flux around single and multiple tufts of hyaline hairs on the thallus of *Fucus vesiculosus*. Flow was a major determinant of DBL thickness, where higher flow decreased DBL thickness and increased O_2_ flux between algal thallus and the surrounding seawater. However, the topography of the DBL varied and did not directly follow the contour of the underlying thallus. Areas around single tufts of hyaline hairs exhibited both increased and decreased DBL thickness as compared to areas over smooth thallus surfaces. Over thallus areas with several hyaline hair tufts, the overall effect was a local increase in the DBL thickness. We also found indications for advective O_2_ transport driven by pressure gradients or vortex-shedding downstream from dense tufts of hyaline hairs alleviating local mass-transfer resistance imposed by thickened DBL. Mass-transfer dynamics around hyaline hair tufts are thus more complex than hitherto assumed and may have important implications for algal physiology and plant-microbe interactions.

## Introduction

Aquatic macrophytes are limited compared to their terrestrial counterparts by the ∼10^4^ slower diffusion and a much lower solubility of gases in water than in air [1, 2]. The efficient exchange of nutrients and gases is further exacerbated by the diffusive boundary layer (DBL) surrounding all submersed surfaces [3]. The thickness and topography of diffusive boundary layers around submerged impermeable objects is affected by flow velocity and surface topography [3]. Higher flow velocities decrease the DBL thickness by exerting a higher shear stress on the viscous sublayer near the surface. The effect of surface roughness on DBL thickness is variable, where angled planes facing the flow generally will have decreased boundary layers, while the DBL downstream of protruding structures will have increased boundary layers. At the same time, the effect of surface roughness on mass transfer across a boundary layer is dichotomous, where a thicker DBL will decrease mass transfer, while surface roughness tends to increase the overall surface area increasing mass transfer [3]. In microsensor-based studies of the DBL, it is important to note that the presence of the microsensor tip in itself can affect the local DBL thickness [4], where flow acceleration around the microsensor shaft will compress the local DBL thickness leading to locally enhanced O_2_ fluxes in the order of 10%. However, this effect is only significant when investigated on smooth surfaces, while a clear effect is apparently undetectable when measured over tufts in e.g. a cyanobacterial mat [5].

It has been estimated that the DBL accounts for up to 90% of the resistance to carbon fixation in freshwater plants [6], and structural-and biochemical regulations to alleviate such mass transfer resistance have evolved across lineages. Some aquatic macrophytes have e.g. developed i) thinner leaves and a reduced cuticle that decreases the diffusion path length to chloroplasts, ii) carbon concentrating mechanisms that increase internal carbon concentration, and iii) the ability to utilize HCO_3_^−^ which constitutes the largest fraction of dissolved inorganic carbon at ocean pH ([7], [8] and references therein). In photosymbiotic corals it has also been proposed that vortical ciliary flow can actively enhance mass transfer between the coral tissue and the surrounding water in stagnant or very low flow regimes with concomitant thick diffusive boundary layers [9].

Hyaline hairs, i.e., colourless, filamentous multicellular structures, are often observed as whitish tufts on the thallus of brown macroalgae in the genus *Fucus*. The hairs originate and are anchored in cryptostomatal cavities on the apical-and mid-regions of the thallus [10]. It is recognized that hyaline hairs aid in the uptake of nutrients [11, 12] e.g. during springtime, when photosynthetic potential is higher due to increased light levels and the need for nutrients apparently triggers growth of hyaline hairs [10].

How hyaline hairs affect solute exchange and nutrient acquisition in *Fucus* is still debated, although a number of suggestions currently exist in the literature suggesting that: i) the hairs increase the algal surface area available for nutrient uptake, albeit this is probably not the major limitation on nutrient uptake [11]; ii) the hyaline hairs might decrease the diffusive boundary layer (DBL) due to turbulence created by the hairs as water flows across them, decreasing the mass resistance imposed by the DBL [11]; iii) the thin cell walls of the hairs relative to the thallus could have less resistance to the passage of ions [13, 14]; iv) the hyaline hairs increase DBL thickness, thereby retaining the products of thallus surface-active enzymes such as extracellular phosphatases and ensuring more efficient nutrient uptake [14].

There can be no doubt, however, that the DBL has great importance for macroalgal growth rates. The mass resistance imposed by the DBL has been correlated with nutrient limitation for the giant kelp *Macrocystis pyrifera* [15], and a considerable spatial variation of the DBL over thallus and cryptostomata of *Fucus vesiculosus* has been observed [16]. However, current knowledge of the DBL characteristics of aquatic plants is largely based on point measurements with O_2_ microsensors [14], while it is known from boundary layer studies in biofilms [17], corals [18–20] and sediments [17, 21–23] that the DBL exhibits a spatio-temporal heterogeneity, which is modulated by both flow velocity and surface topography. Similar studies of DBL topography are very limited in aquatic plant science[24] and the aim of this study was to explore how the DBL thickness and the local O_2_ flux varied spatially over the thallus of *F. vesiculosus* with and without tufts of hyaline hairs. Such first time exploration of the 3D DBL topography was done with O_2_ microsensors mounted in a fully automated motorized micromanipulator system that allowed measurements of DBL transects and grids over the thallus of *F. vesiculosus* (Fig. S1). Our results reveal a complex DBL landscape over the algal thallus, where mass transfer across the DBL apparently can be supplemented by advective processes due to the presence of hyaline hairs.

## Materials and Methods

### Sampling and experimental setup

Specimens of *Fucus vesiculosus* and seawater used in the experimental setup were sampled on the day of usage at <1 m depth at Kronborg, Helsingør, Denmark from May through August. If the influence of a single tuft of hyaline hairs on the DBL was measured, the surface of the thallus surrounding the tuft was carefully shaved off any additional tufts with a scalpel and observation under a dissection microscope to avoid thallus damage. When the influence of multiple tufts was analysed, the thallus was left intact. Prior to measurements, a piece of *F. vesiculosus* thallus with hyaline hairs was fixed on a slab of agar (∼1.5% w/w in seawater) in a small flow chamber creating a defined unidirectional flow of seawater across the thallus surface [16]. The flow chamber was connected via tubing to a submersible water pump in a continuously aerated seawater reservoir tank underneath the flow chamber. Flow velocities were adjusted by restricting water flow to the flow chamber, and flow velocity was determined by collecting water from the flow chamber outlet for one minute, where after the sampled volume per time was divided by the cross sectional area of the flow chamber to obtain an estimate of the mean free flow velocity. All measurements were done with flow velocities of either 1.65 cm s^−1^ or 4.88 cm s^−1^.

The sample was illuminated from above with light from a halogen lamp (Schott KL-2500LCD) equipped with a collimating lens yielding a downwelling photon irradiance (400-700 nm) of ∼350 μmol photons m^−2^ s^−1^, as measured with a quantum irradiance meter (LI-250, LiCor Inc., USA).

### Microsensor measurements

Measurements of O_2_ concentration above the thallus of *F. vesiculosus* were done with Clark type O_2_ microelectrodes (tip diameter 10 μm, OX10, Unisense A/S, Denmark; Revsbech 1989) with a response time of 1-3 seconds and low stirring sensitivity (<2%). The microelectrode was connected to a pA meter (PA2000, Unisense A/S, Denmark) and sensor signals from the pA meter were acquired on a PC via a parallel port-connected A/D converter (ADC-101, Pico Technologies Ltd., England). The O_2_ microsensor was mounted in a custom built micromanipulator setup enabling motorized positioning at defined x, y and z coordinates at ∼1 μm resolution by use of 3 interconnected motorized positioners (VT-80, Micos GmbH, Germany) and controllers (MoCo DC, Micos GmbH, Germany). Data acquisition and positioning was controlled by a custom made software (Volfix) programmed in LabView (National Instruments, Japan).

The O_2_ microelectrode signal was linearly calibrated at experimental temperature (∼17°C) and salinity (S=16) from measurements in air saturated seawater and in seawater made anoxic by addition of sodium dithionite. The O_2_ concentration in air saturated seawater, C_0_, and the molecular diffusion coefficient of O_2_, D_0_, in seawater at experimental temperature and salinity was taken from tabulated values (Unisense A/S, Denmark) as C_0_ = 274 μmol O_2_ L^−1^ and D_0_ = 1.8710^−5^ cm^2^ s^−1^.

### Mapping of diffusive boundary layers

The diffusive boundary layer (DBL) around tufts of hyaline hairs anchored in cryptostomata of *F. vesiculosus* was mapped as 2D transects and 3D grid measurements of O_2_ concentration profiles. The Volfix software enabled us to specify a measuring grid/transect with any number of sampling points in the x, y and z-directions. In this study, the y-direction corresponds to the direction of flow (where negative values indicate distance behind a single tuft of hyaline hair), the x-direction corresponds to the width of the flow chamber, and the z-direction corresponds to the height above the thallus surface (Fig. S1). The approximate height and radius of the hyaline hairs was approximated by manual manipulation of the microelectrode tip relative to the structures as observed under a stereomicroscope (SV6, Zeiss, Germany). Thallus samples were placed in the flow chamber with the length of the thallus oriented along the direction of flow, i.e., the y-direction.

*2D DBL transects*. For 2D transect measurements, the O_2_ microelectrode tip was positioned manually as close to the centre of the selected cryptostomata with hyaline hairs as possible using a dissection microscope for observation; this position was set to y=0 in the Volfix measuring software. The transect measurements started 2 mm upstream (y=2 mm) from the cryptostomata and 1 mm above the hyaline hairs (Point A in Fig. S1D), and ended 4 mm downstream (y=−4 mm). Transects of O_2_ concentration profiles were measured at a lateral resolution of 0.5 mm in the y-direction, with vertical O_2_ concentration profiles done at each transect point in steps of 0.1 mm in the z-direction. All profile measurements started at the same z-position and were finished in the upper thallus layer, where a characteristic jump in the O_2_ concentration, due to the physical impact of the O_2_ microsensor and the solid thallus surface, enabled precise determination of the thallus surface.

*3D DBL grids*. For measuring 3D grids of O_2_ concentration profiles over thallus areas with only one tuft of hyaline hairs, 9 transects covering a 24 mm^2^ sampling grid area around a central tuft of hairs was set up (Fig. S1E) for O_2_ measurements at 0.5 mm lateral resolution (x and y direction) and 0.1 mm vertical resolution (z-direction). Measurements started ∼1 mm above the hyaline hairs (Fig. S1E).

For measurements of 3D grids of O_2_ concentration profiles over larger thallus areas with multiple tuft of hyaline hairs, an oblong 12 mm by 2 mm measuring grid was set up. Oxygen measurements were done at a lateral resolution of 0.5 mm (x and y directions) and a vertical resolution of 0.2 mm (z direction). Tufts of hyaline hair were scattered across the thallus, and the starting point of the grid measurement was set randomly, but with the same starting point for both the 1.65 cm s^−1^ and 4.88 cm s^−1^ measurements.Due to the length of measurements different thallus samples were used for individual experiments.

### Diffusive boundary layer thickness and calculations

There are different ways of determining the effective thickness of the diffusive boundary layer from O_2_ microsensor measurements [17]. The DBL thickness is often found by extrapolating the linear O_2_ gradient in the DBL to the bulk concentration of the free-flow region. The distance from the surface to the intersection of the extrapolated linear gradient and the bulk concentration is denoted the effective diffusive boundary layer thickness, Z_δ_[3]. However, analysing large numbers of O_2_ profiles in this manner is very time consuming, and a somewhat faster determination can be done by defining Z_δ_ as the distance between the surface and the vertical position above the surface where the O_2_ concentration has changed 10% relative to the O_2_ concentration in the bulk water. Estimations of Z_δ_ via this method were found to differ <10% from more precise determinations [17].

The diffusive flux of O_2_ across the DBL, J (in units of nmol O_2_ cm^−2^ s^−1^) was calculated from steady state O_2_ concentration profiles using Fick’s 1^st^ law:

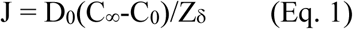

where C_∞_ is the O_2_ concentration in the free-flow region (μmol O_2_ L^−1^ = nmol O_2_ cm^−3^), C_0_ is the O_2_ concentration at the thallus surface (μmol O_2_ L^−1^), Z_δ_ is the effective DBL thickness (cm), and D_0_ is the molecular diffusion coefficient of O_2_ in seawater (cm^2^ s^−1^). To compensate for the uneven surface of thalli, measurements below the thallus surface are not shown on final transects. The depth axis on transects is therefore denoted z’ = z − z_0_, where z is the z-coordinate from the sample data, and z_0_ is the z-coordinate of the thallus surface. The thallus surface position was determined from the intermittent sudden drop in O_2_ concentration when the microsensor pushed against the thallus surface cortex. Maps of O_2_ concentration, Z_δ_, and J were generated from measured transects and grids using data interpolation software (Kriging gridding method using default settings, i.e. Linear Variogram and Point Kriging, Surfer v.8, Golden Software Inc., USA).

### Statistics

Two-way ANOVAs were applied to test differences in the mean Z_δ_ over *F. vesiculosus* between flow rates and light conditions (light/dark). For significant main effects (flow rate and/or light condition) and interaction effects, Tukey’s multiple comparisons post hoc test was applied. Two-way ANOVAs were applied to test differences between mean O_2_ flux values between flow rates and thallus condition (single or multiple tufts). For significant main effects (flow rate and/or thallus condition) and interaction effects, Tukey’s multiple comparisons post hoc test was applied. Statistical analysis was performed using Rstudio (Rstudio version 0.99.491, 2016) with the level of significance set to p <0.05.

## Results

### The DBL around single tufts of hyaline hairs

Isopleths of O_2_ concentration in the water column above the thalli showed a local increase in effective DBL thickness, Z_δ_ associated with the hyaline hair tuft (Fig. 1), with highest Z_δ_ values located downstream from the hyaline hairs, either directly behind the tuft or even within the expanse of the hyaline hairs. In light (350 μmol photons m^−2^ s^−1^), and at a flow of 1.65 cm s^−1^, Z_δ_ reached a maximum thickness of 1.2 mm downstream relative to the tuft at y = −0.5 mm (Fig. S2, S3A), while under a flow of 4.88 cm s^−1^, the maximum Z_δ_ was reduced to 0.6 mm at y = −2.5 mm (Fig. S3A). In darkness, the maximal Z_δ_ values were 0.9 mm and 0.4 mm at y = −2 mm and y = 0 mm under flows of 1.65 and 4.88 cm s^−1^, respectively. The mean Z_δ_ did not change significantly (p<0.05) between measurements in light and darkness (Fig. S3B) under low flow (Z_δ(Light)_ = 0.72 mm, Z_δ(Dark)_ = 0.58 mm; p adj = 0.26) or high flow (Z_δ(Light)_ = 0.36 mm, Z_δ(Dark)_ = 0.18 mm; p adj = 0.13). However, flow velocity had a significant effect on the mean Z_δ_ that was significantly thinner under high flow (4.88 cm s^−1^) as compared to the low flow (1.65 cm s^−1^) (p adj < 0.001 for both main effects).

**Figure 1.**
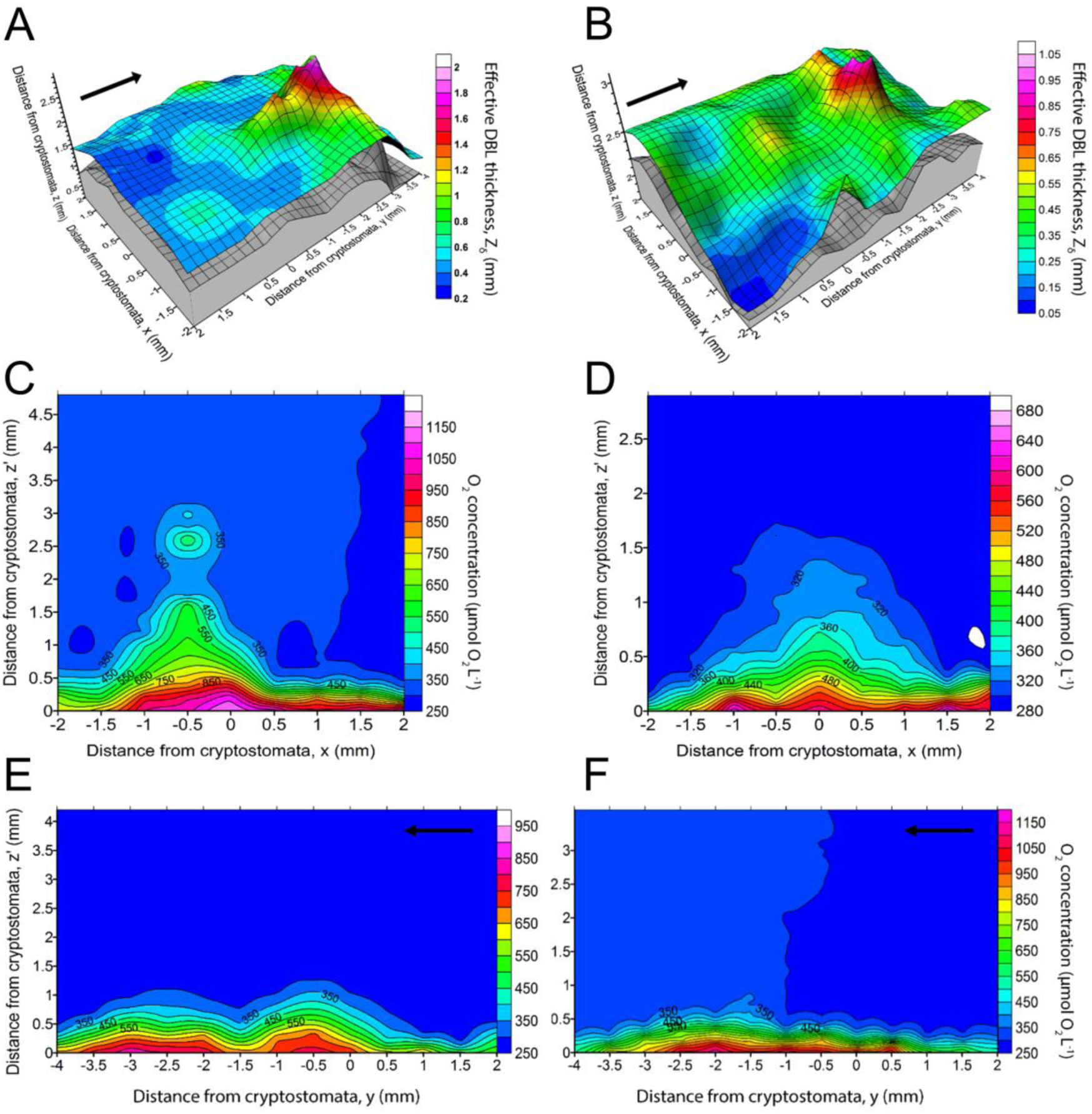
Fine-scale mapping of DBL around a single tuft of hyaline hairs on an illuminated *Fucus vesiculosus* thallus (350 μmol photons m^−2^ s^−1^). (A) and (B) 3D plots of *F. vesiculosus* thallus surface (grey area) and upper extension of DBL (coloured area) around a single tuft of hyaline hairs, at flow velocities of 1.65 (left panels) and 4.88 cm s^−1^ (right panels). Colour bars depict the effective DBL thickness, Z_δ_ (mm), and arrows indicate the direction of flow. (C) and (D) Transects in the x-direction (perpendicular to the flow), at position y=-2 mm from Fig. 1 A,B, respectively, normalized to thallus surface showing the local O_2_ concentration. The zero position (0,0) indicates the position of the cryptostomata. Colour bars denote O_2_ concentration (in μmol O_2_ L^−1^). (E) and (F) Transects of O_2_ concentration (in μmol L^−1^) measured across a single tuft of hyaline hairs in *Fucus vesiculosus* measured at flow velocities of 1.65 (E) and 4.88 cm s^−1^ (F), in light (350 μmol photons m^−2^ s^−1^). The arrows indicate flow direction. The zero position (0,0) indicates the position of the cryptostomata, and transects were adjusted to the thallus surface. Colour bars denote O_2_ concentration (in μmol O_2_ L^−1^).

The hyaline hairs affected Z_δ_ downstream from the hair tuft (Fig. 1A,B) causing a thickening of the DBL that also expanded perpendicular to the flow direction (Fig. 1C,D), reaching a maximum expansion at y = −2 mm at both flows (Z_δMax_= 1.8 mm for 1.65 cm s^−1^ and Z_δMax_=0.9 mm for 4.88 cm s^−1^). Beyond the local peak in boundary layer thickness, the DBL closely followed the contours of the thallus surface topography. Two transects measured at higher resolution along the x-axis at y=-2 mm showed that the increase in Z_δ_ was roughly identical and extended ∼1 mm on both sides of the hyaline hair tuft (Fig. 1C,D). At distances >1 mm away from the local maximum, the DBL thickness approached a lower more homogeneous DBL thickness over thallus areas unaffected by the hair tuft.

Unexpectedly, a transect of O_2_ concentrations above the illuminated thallus at y= −2 mm in the x direction, i.e., perpendicular to the flow, showed a local O_2_ increase followed by a decrease in oxygen concentration that was apparently independent of Z_δ_ (Fig. 1C). A longitudinal transect at x=-0.5 mm along the y-direction, i.e., the flow direction, showed further indications of an apparent local “upwelling” of O_2_ into the transition zone between the DBL and the fully mixed water column protruding downstream from the hyaline hair tuft (Fig. 2). We found such “upwelling” zones most pronounced under low flow located ~2 mm downstream from the centre of the tuft and extending several mm into the water column with O_2_ concentrations reaching up to >2 times air saturation in some cases (Fig. 2A).

**Figure 2.**
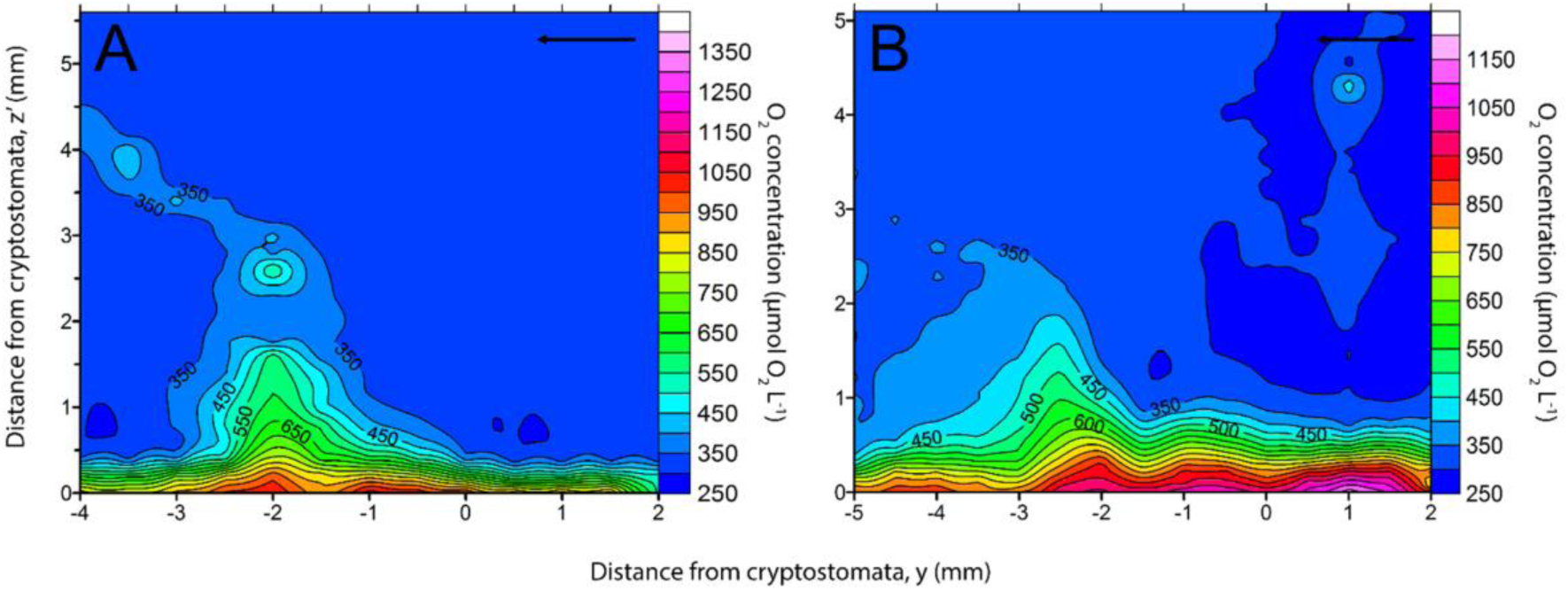
Local transects of O_2_ concentration around single hair tufts from two different measurement series over an illuminated *Fucus vesiculosus* thallus (350 μmol photons m^-2^ s^−1^). (A) shows a transect taken from Fig. 1A at x=-0.5 under a flow velocity of 1.65 cm s^−1^, while (B) was measured similarly as Fig. 1E, also at a flow velocity of 1.65 cm s^−1^. The hair tufts were 2.5-3 mm in diameter and protruded 3-3.5 mm from the thallus. Both transects were adjusted to the thallus surface. The black arrow indicates the flow direction. Colour bars denote O_2_ concentration (in μmol O_2_ L^−1^).

### The DBL around multiple tufts of hyaline hairs

3D grid measurements of O_2_ concentration over a *F. vesiculosus* thallus with several tufts of hyaline hairs spaced at approximately 2-5 mm distance were done to investigate potential combined effects of multiple tufts on the DBL. Such measurements showed that the smooth local thickening of the DBL around a single hyaline tuft relative to the DBL of the smooth thallus was altered in the presence of multiple tufts (Fig. 3). The DBL topography was more heterogeneous with Z_δ_ varying >1 mm reaching a maximum thickness of >2.5 mm under low flow and >1.5 mm under high flow, respectively. The DBL topography was apparently largely determined by the interaction between flow and the tufts of hyaline hairs under low flow, while the smooth thallus surface topography led to local minima in Z_δ_ in-between individual tufts at higher flow velocity (Fig. 3B).

**Figure 3.**
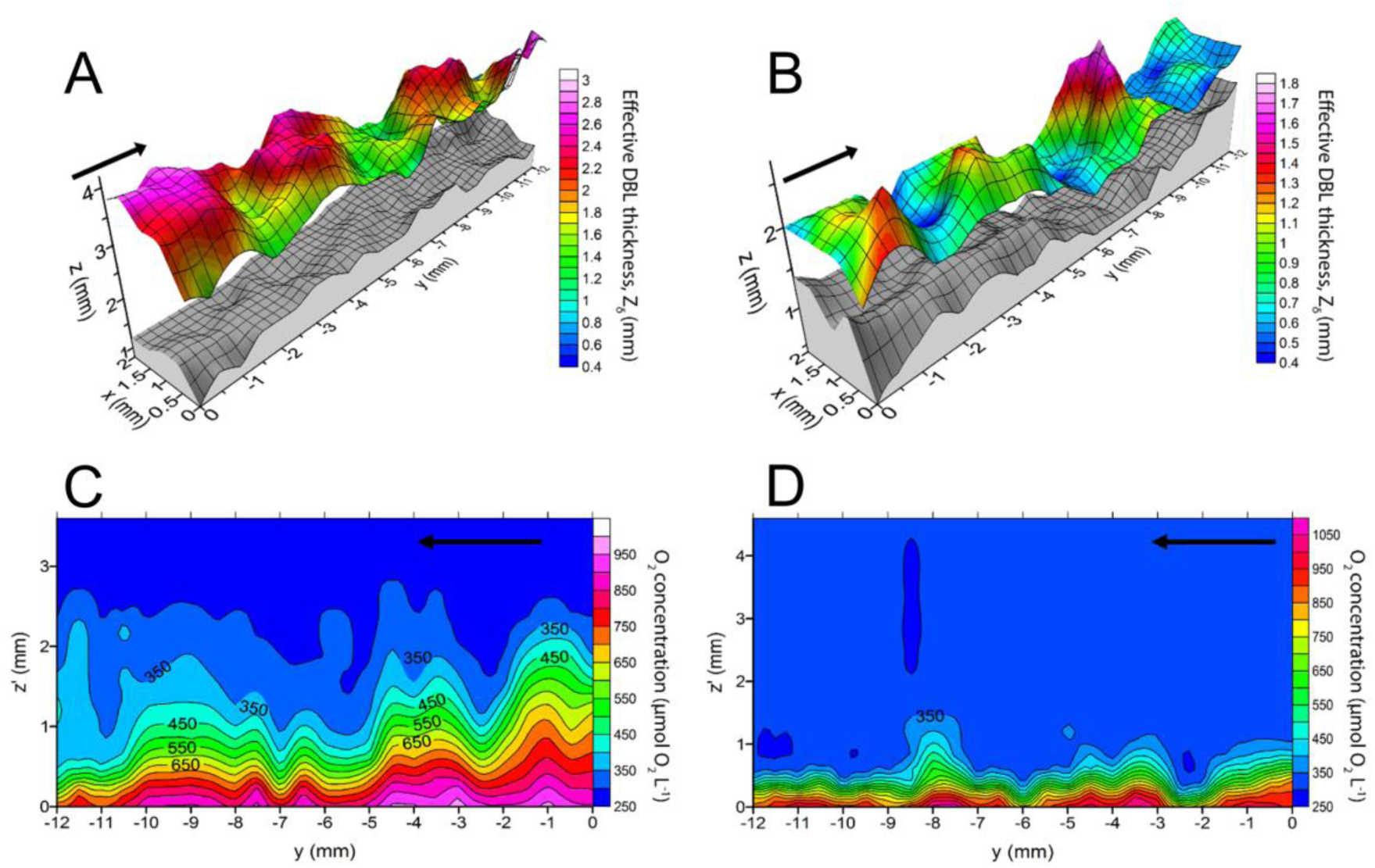
Mapping of DBL over several tufts of hyaline hairs on an illuminated *Fucus vesiculosus* thallus (350 μmol photons m^−2^ s^−1^). (A) and (B) 3D plot of *F. vesiculosus* thallus surface (grey area) and the upper extension of the DBL (coloured area) of multiple tufts of hyaline hairs under a flow velocity 1.65 (left panels) and 4.88 cm s^−1^ (right panels). Colour bars depict the effective DBL thickness, Z_δ_ (mm). (C) and (D) Transects of O_2_ concentration at position x=1 (along the y-axis direction) normalized to thallus surface. Colour bars denote O_2_ concentration (in μmol O_2_ L^−1^).

Transects of O_2_ concentrations at x=1 mm (extracted from the 3D grids in Fig. 3A,B) gave a detailed information on how O_2_ concentration varied over the thallus with distance along the thallus in the flow direction (Fig. 3C,D). In light, the thallus surface O_2_ concentration reached >900 μm both flows, while the O_2_ concentration in the transient zone of the DBL (z=0.7 mm) varied between 350 and 750 μM under low flow and between 300 and 550 μM in high flow, respectively, clearly demonstrating a compression of the boundary layer and more effective O_2_ exchange between thallus and water under higher flow.

However, the thickening of the 300-350 μM O_2_ contour areas e.g. at y=-11 mm and y=-6 mm (Fig. 3C) was due to gradual increasing O_2_ concentrations from the bulk water towards the upper part of the DBL (data not shown). This creates an artefact in the precise determination of Z_δ_ by the method proposed by Jørgensen and Des Marais [17] that will overestimate the local DBL thickness e.g. compared to the local profile in y=-1 mm where a more steady O_2_ increase was measured.

### Diffusive O_2_ fluxes over the *Fucus* thallus with single and multiple tufts

Although inconsistencies were found (e.g. in the area around x = −1.5 mm, y = 1.5mm in Fig. 4A,B), increases in DBL thickness generally correlated with a decrease in O_2_ flux, and the flow-dependent DBL topography strongly affected the O_2_ flux from the illuminated *F. vesiculosus* thallus.

**Figure 4.**
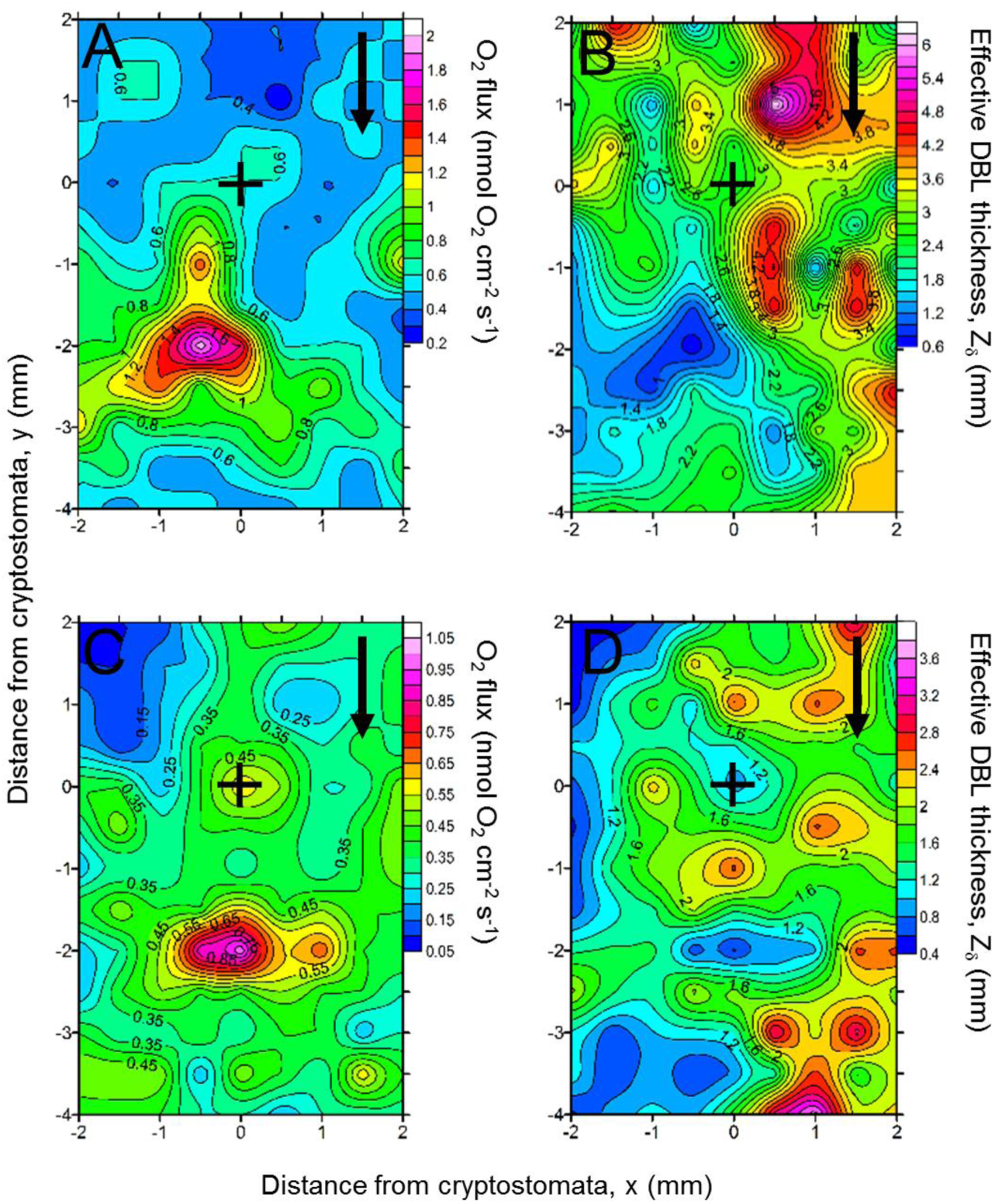
Isopleths of O_2_ flux (in nmol O_2_ cm^−2^ s^−1^) (A) and (C) and effective DBL thickness,Z_δ_(in mm) (B) and (D) measured over an illuminated *Fucus vesiculosus* thallus (350 μmol photons m^−2^ s^−1^) around a single tuft of hyaline hairs at flow velocities of 1.65 cm s^−1^ (A, B) and 4.88 cm s^−1^ (C, D). The hyaline hairs were rooted in the cryptostomata located at the (0,0) coordinate, as indicated by the black cross. Black arrows indicate the flow direction.

Comparing the O_2_ fluxes in transects over the *F. vesiculosus* thallus with single and multiple tufts of hyaline hairs (Fig. 5A,B) showed an increased O_2_ flux just upstream to the position of the hyaline hair tufts independent of the flow velocity. The flux values generally correlated with the DBL thickness and the flux gradually decreased downstream relative to the hair tuft. However, some local variations were seen in areas exhibiting less uniform increases in O_2_ concentration towards the thallus surface.

**Figure 5.**
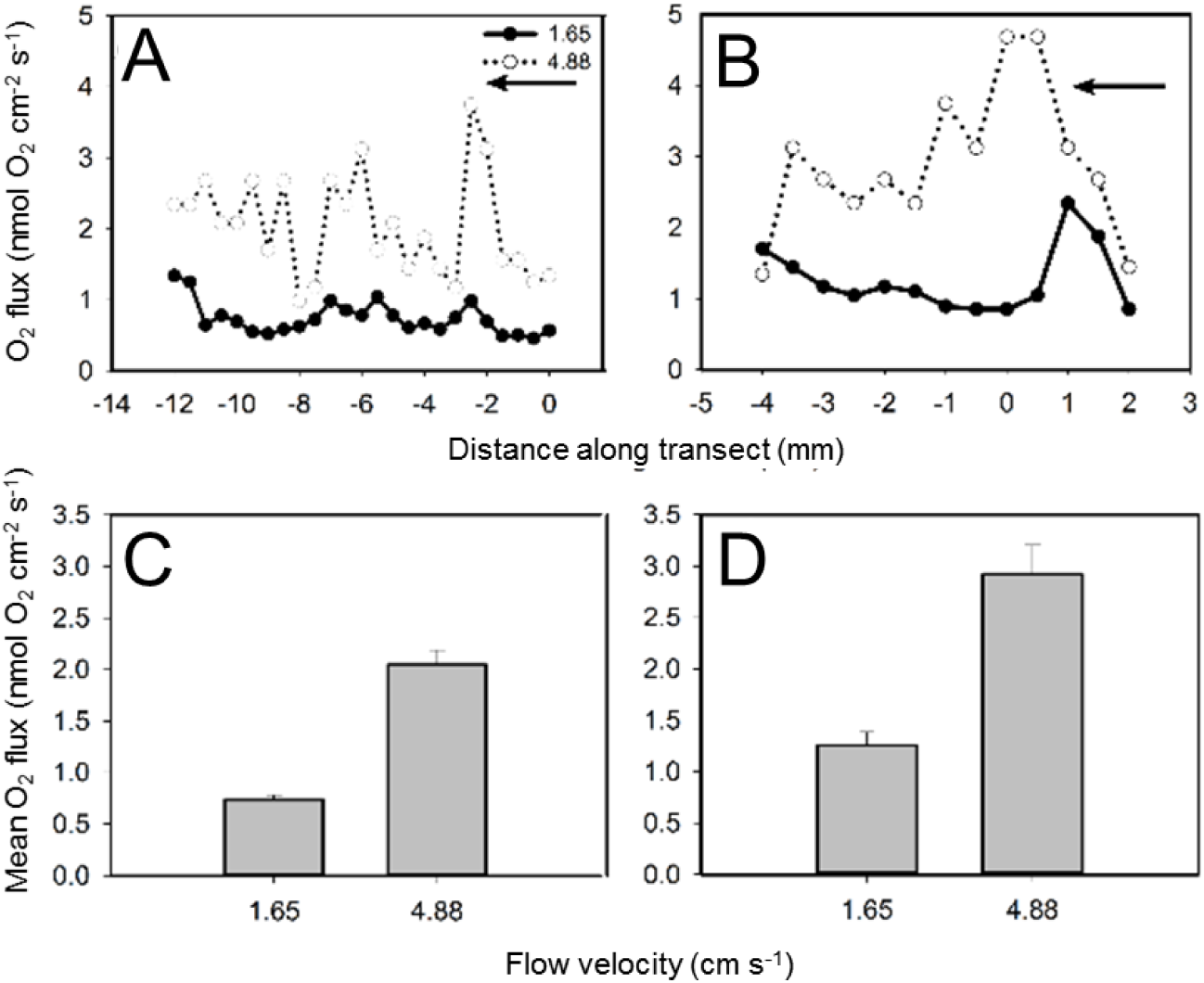
Comparison of O_2_ flux values (in nmol O_2_ cm^−2^ s^−1^) calculated from transects of O_2_ concentration profiles measured over an illuminated (350 μmol photons m^−2^ s^−1^) intact *Fucus vesiculosus* thallus with several tufts of hyaline hairs (A) and a thallus with only a single hair tuft (B) measured under flow velocities of 1.65 cm s^−1^ and 4.88 cm s^−1^. Note the difference in x-scale. The black arrow indicates the flow direction. The individual position of the multiple hairtufts in panel (A) were not mapped and the zero position on the x-axis thus only reflects the starting point of the transect. In panel (B), the zero position indicates the centre of the cryptostomata. Panel (C) and (D) show the average O_2_ flux (±SEM) across (C) the intact thallus and (D) the thallus with a single hair tuft protruding.

The average O_2_ flux calculated from transects over *F. vesiculosus* thalli with single and multiple hyaline hair tufts (Fig. 5C,D) showed that flow was the major determinant of gas exchange between macroalga and the surrounding seawater. Both in measurements over single and multiple hair tufts, the O_2_ flux values were higher in high flow treatments as compared to low flow (p adj <0.001). The O_2_ fluxes measured around a single hair tuft under high flow were higher than the corresponding measurements over multiple tufts treatment (Fig. 5C,D; p adj <0.001). However, the flux values in the multiple tuft treatments were averaged over a two times larger distance (Fig. 5C; 12 mm), and thus include the combined effect of multiple tufts and DBL variation over these, while the values of the single tuft treatments (Fig. 5D; 6mm) only reflect the DBL effects on O_2_ flux immediately downstream from the hair tuft.

## Discussion

In our measurements around single tufts of hyaline hairs, the DBL followed the contour of the smooth thallus surface, except around the hair tufts where a thickening occurred downstream of tufts and perpendicular to the flow direction, with a concomitant decrease in the thickness 1-2 mm away from the hair tufts, depending on the flow-regime.

In measurements over multiple tufts, the smooth thickening of the DBL observed around isolated single tufts was absent. This more dynamic DBL landscape is probably caused by the close vicinity of neighbouring hyaline hair tufts, where the DBL thickness between tufts never reach ‘normal’ conditions over a smooth thallus. Interestingly, this suggests that the effect of multiple hair tufts overall increased the DBL thickness across the thallus. Intuitively, a thin boundary layer would create physical conditions that could better avoid high detrimental O_2_ concentrations and inorganic carbon limitations in light and O_2_ limitation in darkness. So why does *Fucus* spend metabolic energy on production of hyaline hairs? In early work by Raven [11], it was suggested that hyaline hairs aid in nutrient uptake by having a highly decreased diffusion resistance over the plasmalemma compared to the thick algal thallus, and in addition the hairs could protrude through the viscous sublayer and into the mainstream flow with better nutrient access. However, as pointed out by Hurd [14], the thin and flexible hairs are considered unlikely to disrupt the viscous sublayer and create turbulence themselves. Here we show that across a thallus with multiple hair tufts, the overall DBL thickness is increased, which has a functional significance similar to observed DBL effects of epiphytes on submerged macrophytes [24, 25]. The thickening of the boundary layer thickness creates a mass transfer limitation that in light can lead to high O_2_ concentrations [24, 26] potentially inducing photorespiration [27] and limiting the inorganic carbon supply [28] to the thallus. However, such mass transfer limitation would also maintain higher nutrient concentrations due to surface-associated enzyme activity that can aid in the uptake of e.g. phosphorous and other nutrients [14, 29].

A thicker diffusive boundary layer over thalli with tufts of hyaline hairs could also create a niche for epibiotic bacteria, and the presence of bacteria on algal thalli is well known [30–32]. In light of the recently developed ‘holobiont’ concept [32, 33] a physical structure that would keep the algal associated bacteria more protected and provide them with metabolic compounds, could present a competitive advantage. Studies of the role of bacteria in algal life-cycle and metabolism have shown that a strong host specificity of epiphytic bacterial communities exists, possibly shaped by the algal metabolites as the primary selective force [34]. Previous studies have e.g. demonstrated the presence of N_2_ fixing cyanobacteria as part of the algal microbiome, and it has also been shown that native bacteria are required for normal morphological development in some algae [35]. However, the actual distribution and ecological niches of such macroalgae-associated microbes are not well studied. Spilling et al. (2010) found more pronounced O_2_ dynamics, reaching anoxia during darkness, in the cryptostomata cavities of *F. vesiculosus*, wherein the hyaline hairs are anchored. Cryptostomata could thus represent potential niches for bacterial aerobic and anaerobic degradation of organic substrates or O_2_-sensitive N_2_ fixation that warrant further exploration.

In some DBL transects measured on light exposed *Fucus* thalli, we observed areas of enhanced O_2_ concentration detached from the boundary layer (Fig. 2). In a previous study, it was shown that nutrient uptake rates could increase 10-fold when the boundary layer was periodically stripped by passing waves [36]. However, in our case the flow upstream from the tuft was laminar and no waves or DBL stripping occurred. The observed phenomenon of enhanced O_2_ above the DBL could be explained by a combination of factors. As flow is obstructed by a physical object a differential pressure field is created where a local drop in pressure is created around the hyaline hairs due to the locally smaller cross section of unobstructed flow. Such pressure gradient could create a local advective upwelling around the area of low pressure thus affecting the O_2_ transport. The phenomena is well described in e.g. sediment transport [37] and plumes of O_2_ release have also been observed before in coral-reef-associated algae *Chaetomorpha sp.* using planar optodes [38].

In addition, so-called vortex shedding (von Kármán vortex sheets) could also be a factor influencing the observed O_2_ release. Shedding of vortices can occur at certain Reynold numbers at the transition between laminar and turbulent flow when the pressure increases in the direction of the flow, i.e., in the presence of a so-called adverse pressure gradient[39]. In our study, the flow upstream from the hyaline hairs was laminar but a transition to turbulent flow can occur, even at low Reynold numbers, when a certain surface roughness is present and vortex shedding can initiate at Reynold numbers of ∼50 [40]. Using characteristic scales from this study (hyaline hair tuft diameter = 2 mm; free-stream velocity = 1.65 or 4.88 cm s^−1^; fluid density = 1 kg L^−1^ and a dynamic fluid viscosity of 1.08 x 10^−3^ Pa s [41], we calculated Reynold numbers of ∼30-90. Von Kármán vortices have previously been connected to flow patterns on the lee side of plant parts [42] and based on the calculated Reynold numbers the theoretical basis for generation of vortex shedding [40] due to tufts of hyaline hairs is present in our experimental setup.

We thus speculate that a combination of pressure gradient mediated upwelling of O_2_ and vortex shedding (Fig. 6) could explain the observed phenomena in Fig. 2. The local mass transfer related to the presence of hyaline hair tufts on Fucoid macroalgae may thus be more complex. A more detailed investigation of such phenomena was beyond the scope of the present study and would clearly require more detailed characterization of the hydrodynamic regimes over thalli with and without tufts of hyaline hairs.

**Figure 6.**
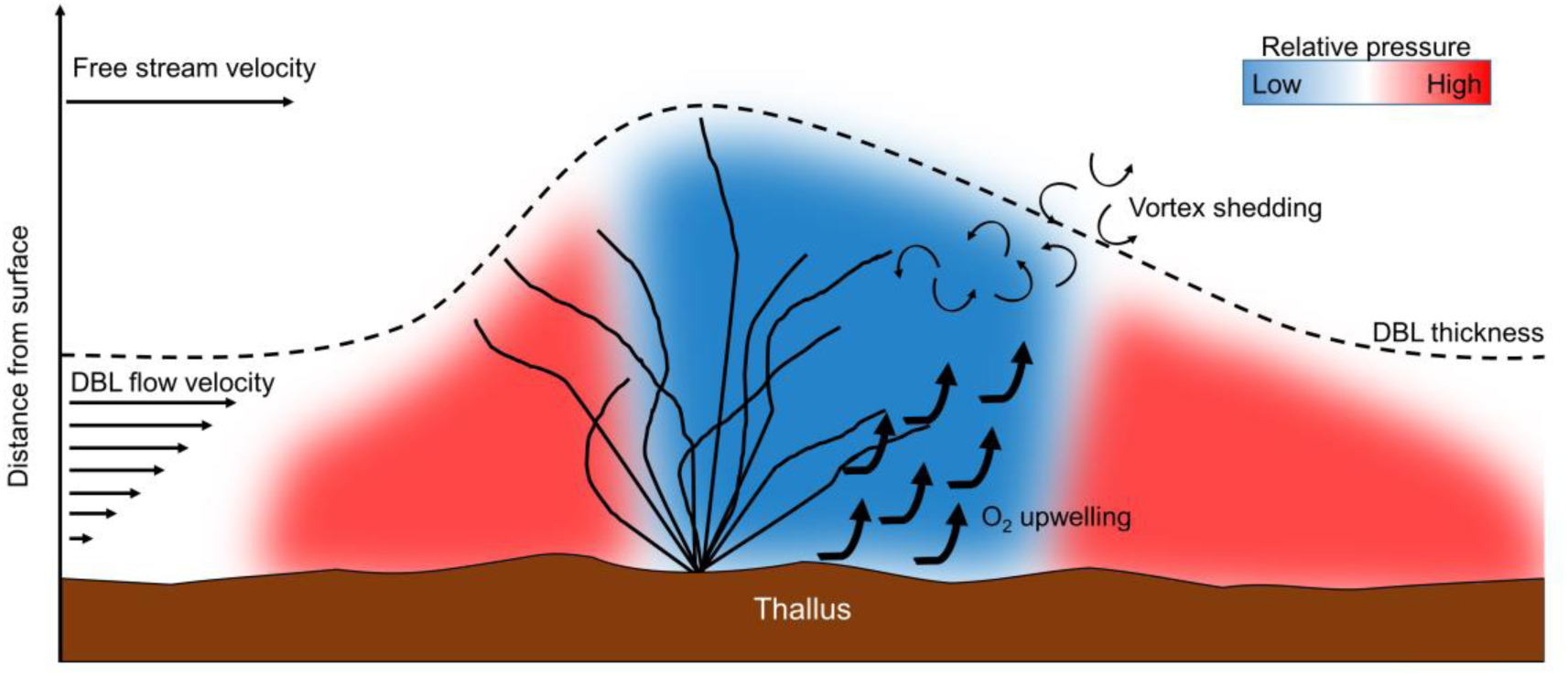
Conceptual drawing showing possible scenarios for the observed upwelling of O_2_ downstream of the hyaline hairs. Flow velocity (straight black lines) decreases from the free stream velocity as the thallus surface is approached through the diffusive boundary layer (DBL). The hyaline hair tuft protruding from cryptostomata alters the DBL thickness locally and creates a differential pressure field (shown in gradient blue and red colours) due to the smaller cross section of unobstructed flow. This creates a flow acceleration in the areas of relative low pressure which could result in advective upwelling. In addition, an adverse pressure gradient is generated downstream from the hair tuft potentially resulting in vortex shedding.

In conclusion, our study of the chemical boundary layer landscape over the thallus of *F. vesiculosus* revealed a strong local effect on the DBL over and around tufts of hyaline hairs anchored in cryptostomata. Single tufts showed a thickening of the DBL downstream and horizontally relative to the thinner DBL over the smooth thallus surface, while areas with multiple tufts exhibited a consistently thickened DBL that may affect gas and nutrient exchanges between the alga and seawater. Furthermore, we also observed more complex solute exchange phenomena that were apparently driven by pressure gradients and/or vortex shedding over the hyaline hair tufts. Altogether, this study demonstrates that interactions between flow and distinct macroalgal surface topography gives rise to local heterogeneity in the chemical landscape and solute exchange that may allow for microenvironmental niches harbouring microbial epiphytes facilitating a diversity of aerobic and anaerobic processes. Further microscale studies of such niches in combination with e.g. microscopy and molecular detection of microbes in relation to hyaline hairs and cryptostomata thus seem an important next step to reveal further insights to the presence and role of the microbiome of Fucoid algae.

## Acknowledgements

This study was supported by grants from the Danish Council for Independent Research | Natural Sciences (MK), and a PhD stipend from the Department of Biology, University of Copenhagen (ML). We thank Roland Thar (Pyro-Science GmbH) for his help in establishing the 3D microsensor measurement setup and software and Erik Trampe for help with photography of Fig. S1C. The authors declare no conflict of interest and no competing financial interest.

## Author contributions

RDN and MK designed the research, RDN and ML performed the research, ML, RDN and MK analysed data, ML wrote the paper with editorial assistance from RDN and MK. All authors gave final approval for publication.

